# Machine learning classification can reduce false positives in structure-based virtual screening

**DOI:** 10.1101/2020.01.10.902411

**Authors:** Yusuf Adeshina, Eric Deeds, John Karanicolas

## Abstract

With the recent explosion in the size of libraries available for screening, virtual screening is positioned to assume a more prominent role in early drug discovery’s search for active chemical matter. Modern virtual screening methods are still, however, plagued with high false positive rates: typically, only about 12% of the top-scoring compounds actually show activity when tested in biochemical assays. We argue that most scoring functions used for this task have been developed with insufficient thoughtfulness into the datasets on which they are trained and tested, leading to overly simplistic models and/or overtraining. These problems are compounded in the literature because none of the studies reporting new scoring methods have validated their model prospectively within the same study. Here, we report a new strategy for building a training dataset (D-COID) that aims to generate highly-compelling decoy complexes that are individually matched to available active complexes. Using this dataset, we train a general-purpose classifier for virtual screening (vScreenML) that is built on the XGBoost framework of gradient-boosted decision trees. In retrospective benchmarks, our new classifier shows outstanding performance relative to other scoring functions. We additionally evaluate the classifier in a prospective context, by screening for new acetylcholinesterase inhibitors. Remarkably, we find that nearly all compounds selected by vScreenML show detectable activity at 50 µM, with 10 of 23 providing greater than 50% inhibition at this concentration. Without any medicinal chemistry optimization, the most potent hit from this initial screen has an IC_50_ of 280 nM, corresponding to a Ki value of 173 nM. These results support using the D-COID strategy for training classifiers in other computational biology tasks, and for vScreenML in virtual screening campaigns against other protein targets. Both D-COID and vScreenML are freely distributed to facilitate such efforts.

## Introduction

Advances in biomedical sciences, driven especially by the advent of next-generation genome sequencing technologies, have enabled discovery of many new potential drug targets [1,2]. Ultimately, however, validating a new candidate target for therapeutic intervention requires development of a chemical probe to explore the consequences of pharmacological manipulation of this target [3]. In recent years this step has typically been carried out by using high-throughput screening (HTS) [4] as a starting point for subsequent medicinal chemistry optimization; with improvements in automation, it has become feasible to screen libraries that exceed a million compounds [5].

More recently, however, sets of robust chemical transformations from available building blocks have been used to enumerate huge libraries of compounds that are readily accessible but never before synthesized [6-9]. These libraries can comprise billions of compounds, and thus remain far beyond the scale accessible to even the most ambitious HTS campaign. This expansion of chemical space in which to search, along with the high cost of setting up and implementing an HTS screen, has increasingly driven the use of complementary computational approaches.

In broad terms, virtual screening approaches can be categorized into two classes: ligand-based screens and structure-based screens [10-12]. Ligand-based screening starts from the (2D or 3D) structure of one or more already-known ligands, and then searches a chemical library for examples that are similar (in either a 2D or a 3D sense). In contrast, structure-based screening does not require *a priori* knowledge of any ligands that bind to the target protein: instead, it involves sequentially docking each member of the chemical library against the three-dimensional structure of the target protein (receptor) and using a scoring function to evaluate the “quality” of each modeled protein-ligand complex. The scoring function is intuitively meant to serve as a proxy for the expected strength of a given protein-ligand complex (i.e. its binding affinity) [13], and is typically built upon either a physics-based force-field [13-17], an empirical function [18-22], or a set of knowledge-based terms [23-28].

After docking, the scoring function is used to select the most promising compounds for experimental characterization; at this stage the accuracy of the scoring function is of paramount importance, and represents the primary determinant of success or failure in structure-based screening [29]. A snapshot of the field was captured by a review summarizing successful outcomes from 54 virtual screening campaigns against diverse protein targets [12]; for the most part, all groups screened the same 3-4 million compounds from ZINC [30,8]. Excluding GPCR’s and artificial cavities designed into protein cores, the median values across the set reveal that an expert in the field – using their own preferred methods of choice, which can include various post-docking filters and human visual inspection (“expert hit-picking”) – can expect about 12% of their predicted compounds to show activity. That said, the hit rate can also be higher in cases where the composition of the screening library is restricted to compounds containing a functional group with natural affinity for the target site (certain well-explored enzyme active sites). Conversely, the hit rate is typically lower when the scoring function is applied without additional filters or human intervention [12]. The median value of the most potent hit from each of the collected campaigns had K_d_ or K_i_ value of ∼3 µM, although this latter result is strongly impacted by the fact that some of these K_d_ or K_i_ values are from custom compounds subsequently optimized via medicinal chemistry, rather than from the initial screening hit.

Despite extensive efforts, the reasons for which active compounds are only identified at a relatively low rate are not quite clear. In addition to factors not evident from the structure of the modeled complex (compound solubility, incorrectly modeled protonation/tautomerization states of the ligand, etc.), we and others have hypothesized that the current bounds of performance may be attributable to limitations in traditional scoring functions [31,32]: these may include inadequate parametrization of individual energy terms, exclusion of potentially important terms, and also failure to consider potential non-linear interactions between terms. For these reasons, machine learning techniques may be especially well-suited for developing scoring functions that will provide a dramatic improvement in the ability to identify active compounds without human expert intervention. However, while machine learning may offer the potential to improve on the high false positive rate of current scoring function, further analysis has revealed that many methods to date reporting promising results in artificial benchmark experiments may have inadvertently overfit models to the training data [33]: this can be a subtle effect of information leakage, occurring when the validation/testing data are not truly non-redundant from the training data. Other studies have shown that apparently impressive performance from deep learning methods can result from detecting systematic differences in the chemical properties of active versus decoy compounds [34]. Either of these artifacts inflates expectations based on benchmark performance, but ultimately leads to non-transferrable and disappointing outcomes when the methods are tested in subsequent prospective evaluations [35-38].

Here, we report the development of a dataset aimed to promote training of a machine learning model designed to be maximally useful in real-world (prospective) virtual screening applications. To build this dataset, we compile a set of “compelling” decoy complexes: a set that mimics representative compounds that might otherwise move forward to experimental testing if generated in the course of a typical virtual screening pipeline. We then use this dataset to train a machine learning classifier to distinguish active complexes from these compelling decoys, with the rationale that this is precisely the step at which standard scoring functions must be augmented. Finally, we apply this model in a *prospective* experiment, by screening against a typical enzyme target (acetylcholinesterase) and testing the top-scoring compounds in a biochemical (wet lab) assay for inhibition of protein activity.

## Results

### Developing a challenging training set

Machine learning methods at varying levels of sophistication have long been considered in the context of structure-based virtual screening [39,31,32,40-46,29,47-54]. The vast majority of such studies sought to train a regression model that would recapitulate the binding affinities of known complexes, and thus provide a natural and intuitive replacement for traditional scoring functions [31,32,41-46,29,47,49,51-54]. The downside of such a strategy, however, is that the resulting models are not ever exposed to any inactive complexes in the course of training: this is especially important in the context of docked complexes arising from virtual screening, where most compounds in the library are presumably inactive. We instead anticipated that a binary classifier would prove more appropriate for distinguishing active versus inactive compounds, and that training would prove most effective if decoy complexes closely reflected types of complexes that would be encountered during real applications.

Building first our set of active complexes, we drew examples from available crystal structures in the Protein Data Bank (PDB). Others have used collections of active compounds for which the structure of the complex is not known, and docked these to obtain a considerably larger set of active complexes [48,49]. The downside of this approach, however, is that mis-docked examples (which may be numerous) are labeled as active during training; this is problematic because mis-docked models do not have appropriate interactions with the protein target that would lead to engagement, and thus should be marked as inactive by the classifier. While restricting examples of active complexes to those available in the PDB drastically limits the number available for training, this strategy ensures that the resulting model will evaluate complexes on the basis of the protein-ligand interactions provided.

Our primary consideration in compiling active compounds for the training set was that the scope of examples should match as closely as possible those anticipated to be encountered when the model is deployed. Training the model on an overly restrictive set of examples would limit its utility (since many cases will be “out of distribution”), whereas training too broadly might limit the resulting model’s performance. Accordingly, we sought to train the model on precisely the type of scenarios that match its intended application. We therefore further filtered the set of active compounds from the PDB to include only ligands that adhere to the same physicochemical properties required for inclusion in our compound library for real screening applications (see *Methods*). This led to a collection of 1383 active complexes, which were then subjected to energy minimization: this prevented us from inadvertently training a model that simply distinguished between crystal structures and models produced by virtual screening.

Turning next to the set of decoy complexes, our primary consideration in compiling the training set was that the decoy complexes should be as “compelling” as possible. If the decoy complexes can be distinguished from the active complexes in some trivial way – if they frequently have steric clashes, for example, or they are systematically under-packed, or they do not contain intermolecular hydrogen bonds – then the classifier can simply use these obvious differences to readily distinguish active versus inactive compounds. In addition to making compelling decoys, the proportion of decoys-to-actives also has a significant effect on the performance of machine learning trained model [55]. In order to achieve a nearly balanced training set, we aimed to include only small number of (very challenging) decoy complexes.

For each active complex, we first used the DUD-E server [56] to identify fifty compounds with physicochemical properties matched to the active compound but completely unrelated chemical structure: this provided a set of compounds compatible in very broad terms for the corresponding protein’s active site, and also ensured that the decoy compounds would not have systematic differences from the active compounds. We then built low-energy conformations of each candidate decoy compound, and screened these against the three-dimensional structure of the active compound using ROCS [57]. From among the fifty candidates, we selected those that best matched the overall shape and charge distribution of the active ligand. Using the structural alignment of the decoy compound onto the active compound, we placed the decoy into the protein’s active site, and carried out the same energy minimization that was applied to the active complexes (**Figure 1a**).

**Figure 1:**
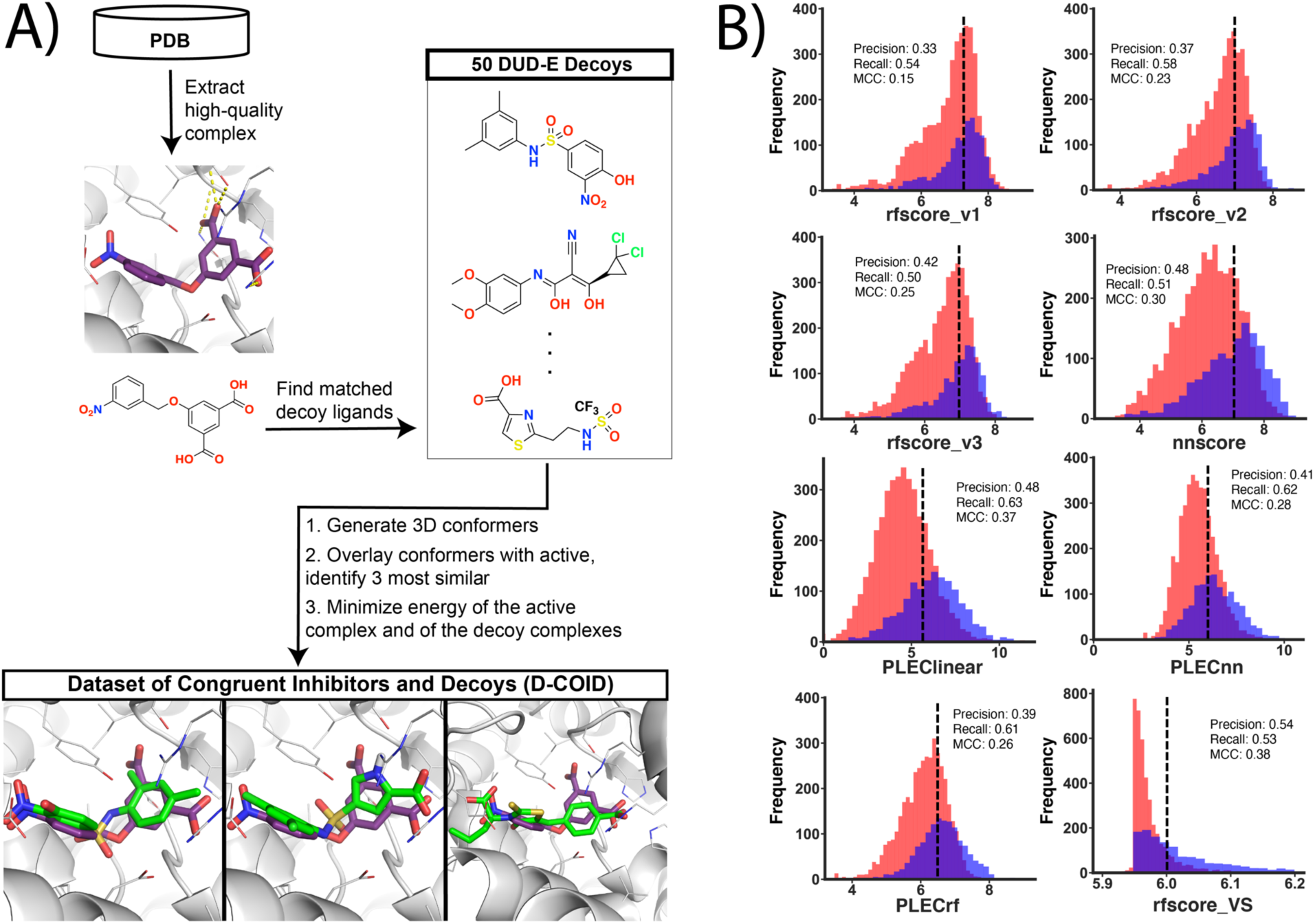
Developing a challenging training set (D-COID). **(A)** Active complexes were assembled from the PDB by filtering for ligands that match those reflected in a screening library. For each active complex, 50 physicochemically-matched compounds were selected and overlaid onto the active compounds; the three most similar compounds on the basis of overall shape and electrostatic similarity were aligned into the protein active site, and used as decoy complexes. This strategy mimics the selection of candidate (active) compounds in a realistic pharmacophore-based screening pipeline, and thus generates highly compelling decoy complexes for training/testing. **(B)** Modern scoring functions cannot distinguish active complexes from decoys in this set. Overlaid histograms are presented for scores obtained using various scoring functions when applied to active complexes (*blue*) and decoy complexes (*red*) in D-COID. For all eight methods tested, the distribution of scores assigned to active complexes strongly overlaps with the distribution of scores assigned to decoy complexes. From each model’s continuous scores, 10-fold cross validation was used to obtain the classification cutoff that maximizes Matthews correlation coefficient (MCC) on each subset of the data. These cutoffs were used in calculating the precision/recall/MCC for each method. The mean of these 10 threshold values is reported with each plot.

We note that the protocol used here to build the decoy complexes doubles as an entirely reasonable approach for ligand-based (pharmacophoric) virtual screening: indeed, ROCS is typically applied to identify compounds with matched three-dimensional properties to a given template, with the expectation that the hits will themselves be active [58-60]. Thus, the unique strategy motivating construction of our training set is in essence a form of adversarial machine learning: we intentionally seek to build decoys that we anticipate would be mis-classified by most models. We named this dataset **D-COID** (**D**ataset of **CO**ngruent **I**nhibitors and **D**ecoys), and have made it publicly-available for others to use freely (see *Methods*).

To confirm that this decoy-generation strategy indeed led to a challenging classification problem, we applied some of the top reported scoring functions in the literature to distinguish between active and decoy complexes in the D-COID set. For all eight methods tested (nnscore [32], RF-Score v1 [31], RF-Score v2 [44], RF-Score v3 [29], PLEClinear [53], PLECnn [53], PLECrf [53], and RF-Score-VS [49]), we found that the distribution of scores assigned to active complexes was strongly overlapping with those of the decoy complexes (**Figure 1b**), indicating that these models showed very little discriminatory power when applied to this set.

Typical scoring functions report a continuous value, because they intend to capture the strength of the protein-ligand interaction. In order to use the scoring function for classification, one must define a threshold value at which complexes are predicted to be either active or inactive. To avoid over-estimating performance by selecting the threshold with knowledge of the test set, we carried out 10-fold cross validation to determine appropriate threshold. In particular, we used 90% of the dataset to define the threshold that maximized the Matthews correlation coefficient (MCC), then applied this threshold to assign each complex in the unseen 10% as active/inactive. Using this unbiased thresholding measure to assign each complex in the D-COID set, we found the Matthews correlation coefficient (MCC) for best-performing scoring function in this experiment to be only 0.39.

### A new classifier for identifying active complexes: vScreenML

Having developed a relevant and challenging training set, we next sought to develop a machine learning model that could discriminate between active and decoy complexes in this set. It has been pointed out in the past that machine learning models built exclusively upon protein-ligand element-element distance counts can yield apparently impressive performance in certain benchmarks without proving useful beyond these [35]. To avoid this pitfall, we used as our starting point the Rosetta energy function [61]: a classical linear combination of traditional (physics-based) molecular mechanics energy terms, alongside empirical terms added so that distributions of atomic arrangements would quantitatively mimic those observed in the PDB [62]. While we acknowledge that the Rosetta energy function is not commonly used for virtual screening, this is primarily because it is too slow to be applied for docking large compound libraries: in one recent benchmark for classification of active versus decoy complexes [63], the Rosetta energy function showed equivalent performance as the popular FRED Chemgauss4 scoring function [64].

At the outset, we found that applying Rosetta to the D-COID set did not yield results notably different than in our previous experiment (**Figure 2a**), and indeed this was confirmed quantitatively through the Matthews correlation coefficient (0.39). Next, we used 10-fold cross validation to re-weight the terms in this scoring function for improved performance in this D-COID classification task using a perceptron [65,66] to maintain the linear functional form of the Rosetta energy function: this resulted in a modest improvement in the apparent separation of scores (**Figure 2b**), but a notable improvement in MCC (0.53). This observation is unsurprising, because the Rosetta energy function is primarily optimized for proteins rather than protein-ligand complexes, and re-training its component energies for a specific task will naturally lead to improved performance for that task. For precisely this reason, historically a separate linearly re-weighted version of the default Rosetta energy function has been used when modeling protein-ligand complexes [67] or when re-ranking complexes from virtual screening [63].

**Figure 2:**
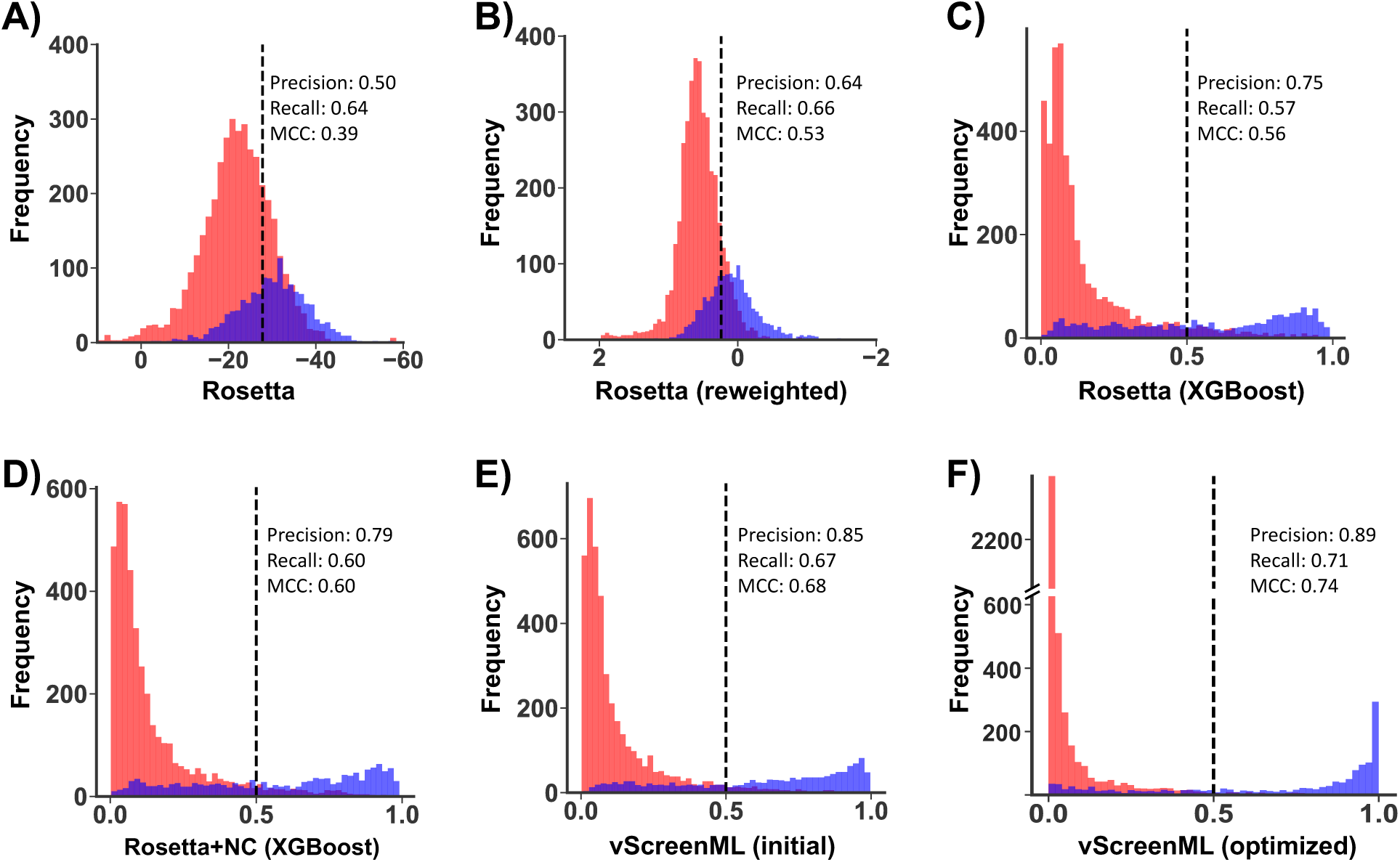
Development of vScreenML. Overlaid histograms are presented for scores obtained when scoring active complexes (*blue*) and decoy complexes (*red*) from D-COID. Scoring functions used were: **(A)** Default Rosetta energy function, **(B)** Linearly-reweighted Rosetta energy terms, **(C)** Rosetta energy terms combined via XGBoost, **(D)** Rosetta energy terms plus structural assessments, **(E)** Rosetta terms plus additional diverse descriptors (non-optimized vScreenML), **(F)** vScreenML after hyperparameter tuning. Over the course of this sequence, the overlap between the active and decoy complexes is progressively reduced and MCC systematically increases. For the first two panels, 10-fold cross validation was used to obtain the classification cutoff that maximizes Matthews correlation coefficient (MCC) on each subset of the data. These cutoffs were used in calculating precision/recall/MCC, and the mean of these 10 threshold values is reported. Because the remaining panels each report results from classification models, their thresholds are fixed at 0.5.

Next, we explored the performance of models that move beyond linear combinations of these energy terms, and instead use these component energies as the basis for building decision trees. Using the XGBoost framework (an implementation of gradient-boosted decision trees), we observed notable separation of the scores assigned to active/decoy complexes (**Figure 2c**), along with a slight increase in MCC (0.56).

To complement the existing terms in the Rosetta energy function, we next added a series of structural quality assessments calculated by Rosetta that are not included in the energy function (**Figure S1**); inclusion of these terms yielded a model with further improved discriminatory power (**Figure 2d**). Inspired by this improvement, we then incorporated additional structural features aiming to capture more sophisticated chemistry than that encoded in Rosetta’s simple energy function, specifically from RF-Score [31] (features that count the occurrence of specific pairwise intermolecular contacts), from BINANA [68] (analysis of intermolecular contacts), from ChemAxon [69] (ligand-specific molecular descriptors), and from Szybki [70] (a term intended to capture ligand conformational entropy lost upon binding). We proceeded to train a model using this collection of features, which we denote “vScreenML”, and were pleased to discover that these again increased the separation between scores assigned to active and decoy complexes (**Figure 2e**). Finally, we used hyperparameter tuning to optimize development of the model (**Figure S2**), and accordingly developed a model that provided nearly complete separation of active and decoy complexes (**Figure 2f**) and unprecedented MCC for this challenging task (0.74). We have made this model publicly-available for others to use freely (see *Methods*).

**Figure S1.**
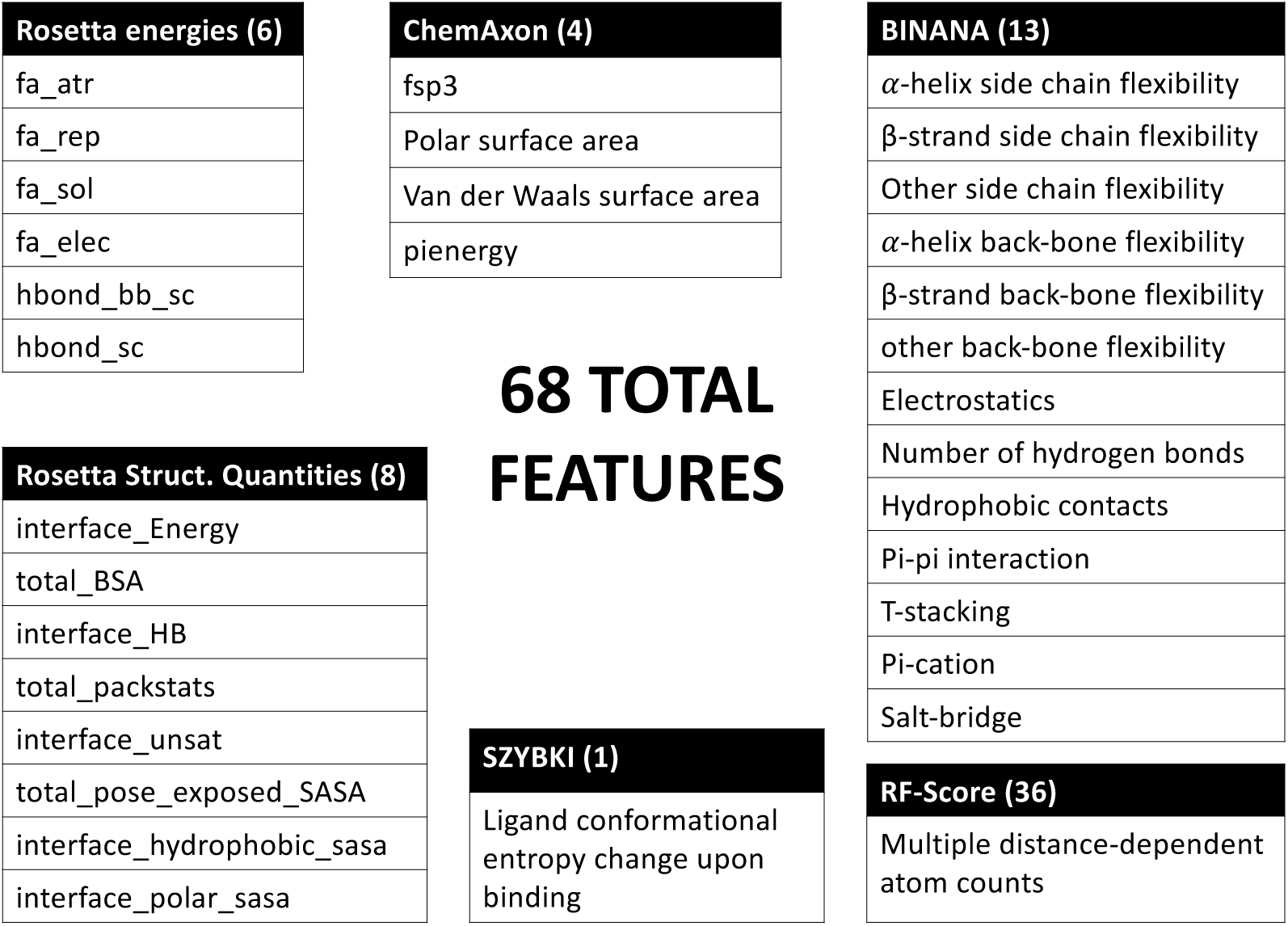
Features incorporated into vScreenML. These features derive from six sources: Rosetta energy terms, Rosetta structural quantifiers, RF-Score’s rfscore_v1 features, BINANA’s analysis of intermolecular contacts, ChemAxon’s cxcalc features, OpenEye’s SZYBKI conformational entropy term.

**Figure S2.**
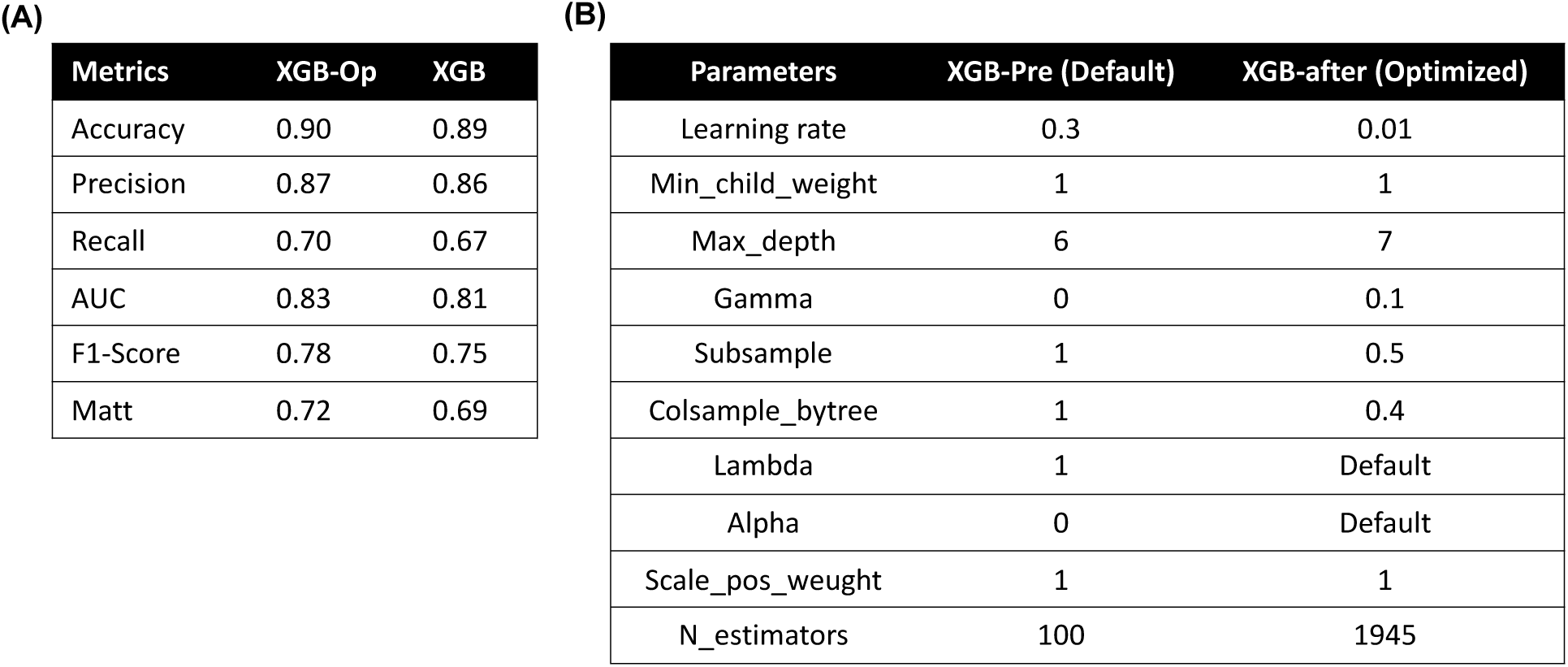
Hyperparameter tuning of vScreenML. **(A)** Performance comparison of the optimized and non-optimized vScreenML models. **(B)** XGBoost parameters before and after optimization.

Through the course of developing of this model, we transitioned from a linear combination of six Rosetta features with clear physical basis, to a collection of 68 diverse and likely non-orthogonal features connected through a more complex underlying model (**Figure S1**). Using the complete set of features that comprise vScreenML, we tested alternate machine learning frameworks, leading us to discover that a different implementation of gradient-boosted decision trees yielded essentially identical performance, and other models built upon decision trees were only slightly worse. By contrast, other models that are not built on decision trees did not provide comparable performance (**Figure S3a**). Importantly, we note that this model has been trained to distinguish actives from decoy complexes in a context where both have been subjected to energy minimization using the Rosetta energy function: the same optimized model is not necessarily expected to recognize actives successfully if they have not been prepared this way (e.g. crystal structures).

**Figure S3.**
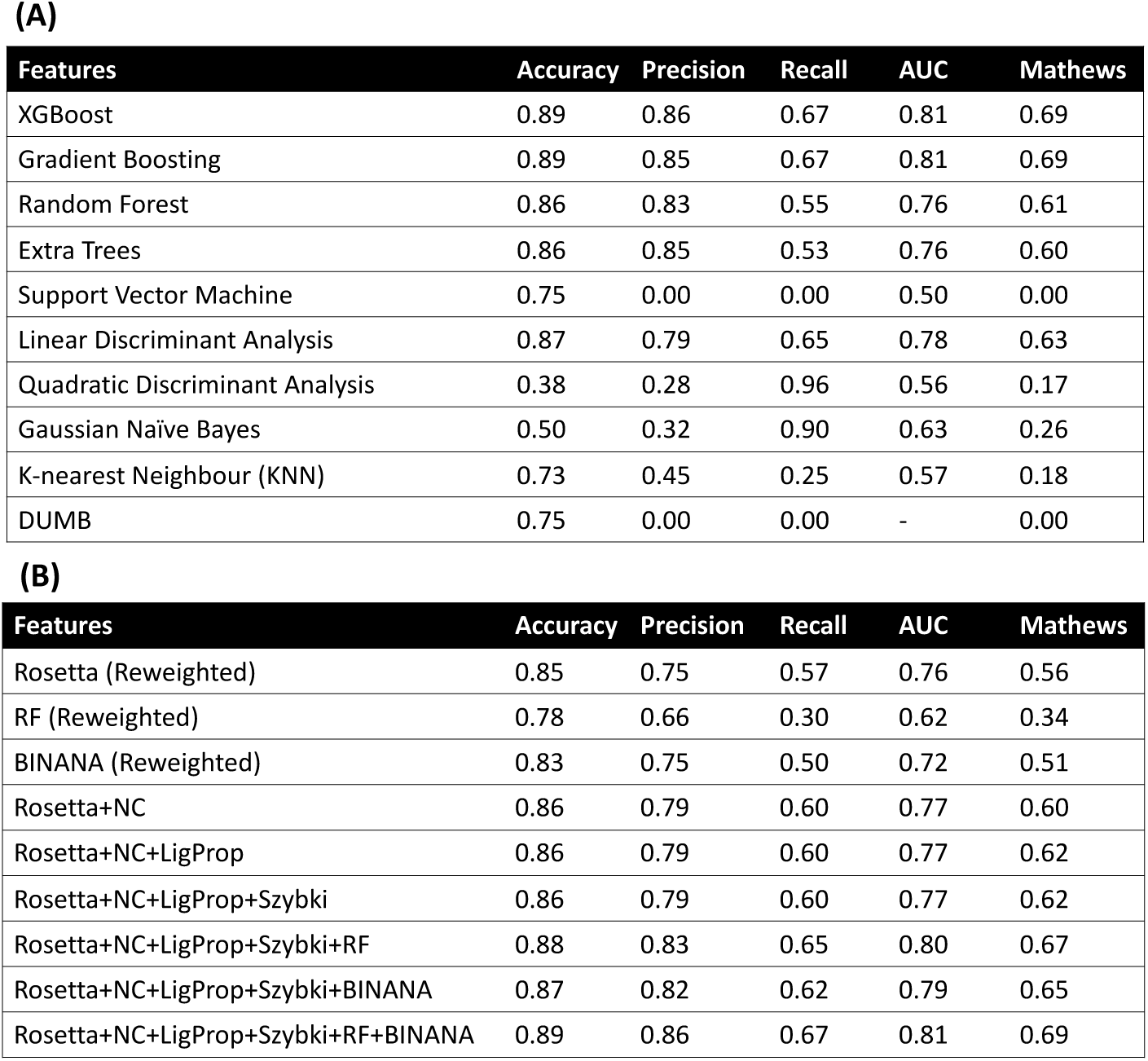
Performance of alternate models. **(A)** Using the complete vScreenML feature set, alternate frameworks are used for building the model. **(B)** Examination of models in which a set of features from a given origin is removed en-masse; all models are trained using XGBoost.

To evaluate the contributions of each part of our feature set, we next removed one at a time all features from a given origin, and explored how the lack of these features would affect performance (**Figure S3b**). This experiment showed that only a very small deterioration in performance was observed when either the RF-Score or BINANA features were removed, but removing both had a large impact; this is unsurprising, given the fact that many of the features in these sets are correlated. Further, removal of SZYBKI’s conformational entropy term had no impact on the model’s performance, suggesting either that the change in ligand conformational entropy as described by SZYBKI does not help distinguish active versus decoy complexes in this dataset, or that this effect is already captured through some combination of other features. In principle, features that are unnecessary (either because they are correlated with other features or because they do not help in classification) should be removed to better avoid the risk of overtraining. In this case, however, we because XGBoost is not particularly susceptible to overtraining and our feature set remains relatively small in comparison to our training set, we elected to instead test our model immediately in orthogonal benchmarks to evaluate potential overtraining.

### Benchmarking vScreenML using independent test sets

The DEKOIS project (currently at version 2.0) [71,72] is intended to provide a “demanding” evaluation set for testing virtual screening methods. Acknowledging that a wide variety of factors make some protein targets easier to model than others, this set includes 81 different proteins with available crystal structures. For each protein, a custom library is provided that contains 40 active compounds and 1200 decoys: thus, about 3.2% of each library is active. The crystal structures of active complexes are not provided (and indeed, most have not yet been experimentally determined). To evaluate performance of a new scoring function, one typically ranks all 1240 compounds for a given protein and selects the top-scoring 12; the enrichment factor for this subset of the library (EF-1%) corresponds to the ratio of the percent of active compounds among the selected 12 to the ratio of active compounds in the original library. Scoring perfectly for a given protein in this set would mean ranking 12 active compounds before all 1200 of the decoys: this would correspond to EF-1% = 1.00/0.032 = 31. Conversely, a method that randomly selects compounds from the library would (on average) select active compounds 3.2% of the time, and thus yield an EF-1% of 1.

Among the 81 proteins in the DEKOIS set, we noted that some were included in our training set as well. To avoid any potential information leakage that might overestimate the performance we could expect in future applications [33], we completely removed these testcases. This left a set of 23 protein targets, each of which vScreenML had never seen before. For each protein, we docked each compound in the corresponding library to the active site (see *Methods*); we note that this unavoidable step could artificially deflate the apparent performance of vScreenML or other models tested, since a mis-docked active compound should have no basis for being identified as active. Some of the compounds in the DEKOIS set could not be suitably modeled in all parts of our pipeline, and were therefore removed; each of the 23 proteins considered ultimately was used to generate 30-40 active complexes and 800-1200 decoy complexes. Each of these complexes (both actives and decoys) were then subjected to energy minimization using the Rosetta: as noted earlier, vScreenML should only be applied in the context of Rosetta-minimized structures. Along with vScreenML, we used eight other machine learning scoring functions were then used to rank the docked-and-minimized models: nnscore [32], PLECnn [53], PLECrf [53], PLEClinear [53], RF-Score v1 [31], RF-Score v2 [44], RF-Score v3 [29] and RF-Score-VS [49]. We additionally included the (default) Rosetta energy function in this benchmark [61].

To compare performance between methods, we plot EF-1% using one method (for each of the 23 protein targets) as a function of EF-1% using the other method (**Figure 3a**). As plotted here, points below the diagonal are specific protein targets for which vScreenML outperformed the alternate method (higher EF-1% for this protein target). The importance of training on both actives and decoys for this task is immediately apparent in these comparisons, by comparing for example vScreenML against PLECnn (a neural network representing the current state-of-the-art among models trained exclusively on active complexes). For the 23 targets in this experiment, PLECnn out-performs vScreenML in 3 cases (points above the diagonal), whereas vScreenML proves superior in 12 cases (the other 8 cases were ties).

**Figure 3:**
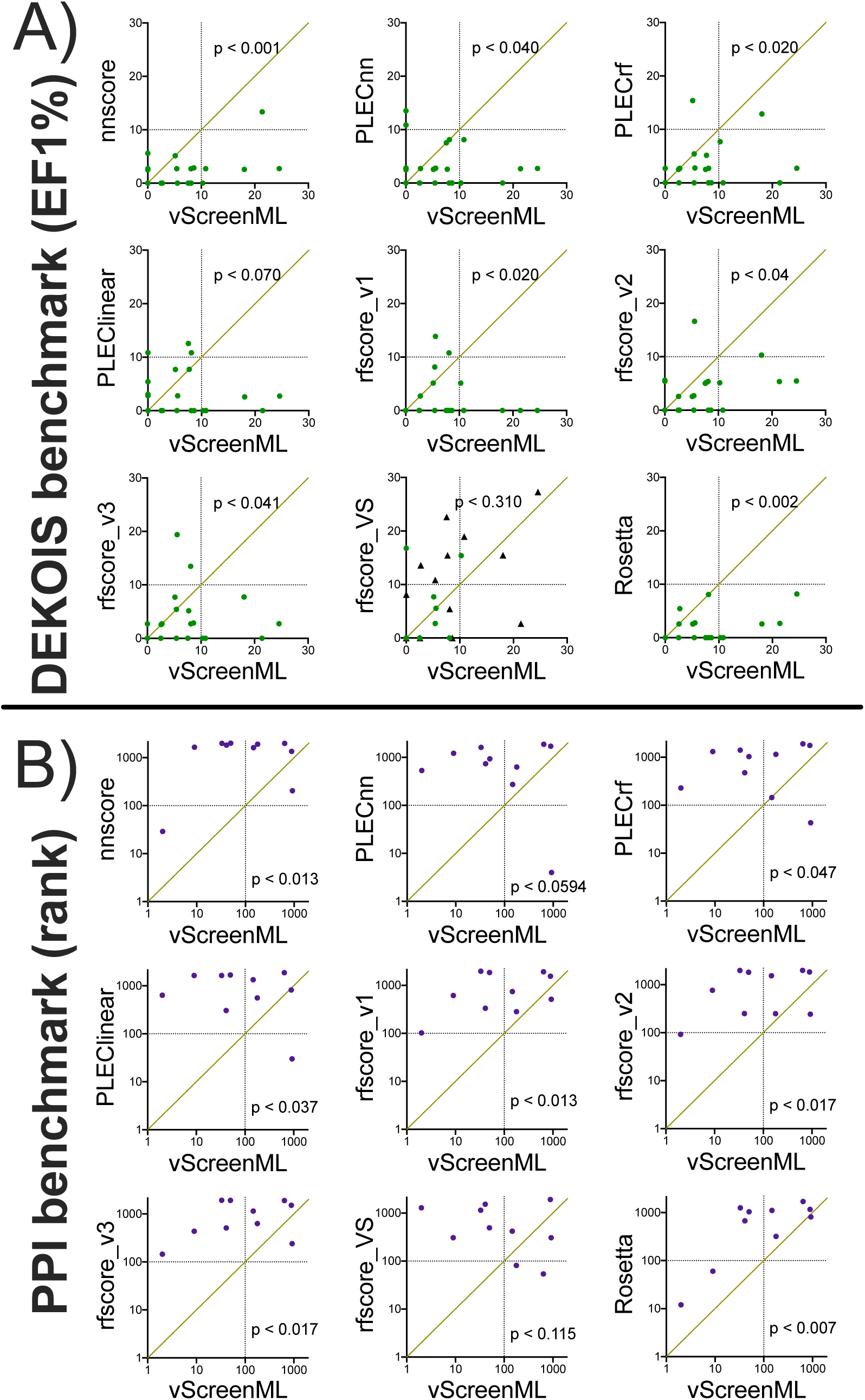
Comparing vScreenML to other scoring functions using two independent virtual screening benchmarks. Each benchmark is comprised of multiple protein targets, corresponding to points on these plots. **(A)** DEKOIS benchmark, comprised of 23 protein targets. For each target (individual dots), 30-40 active complexes and 800-1200 decoy complexes are provided. For a given target, each scoring is used to rank the set of complexes. For a given scoring function, the number of active complexes in the top 1% of all complexes is used to calculate the enrichment of actives relative to randomly selecting complexes; thus, *higher* numbers indicate better performance). When comparing vScreenML against another method, a point *below* the diagonal indicates superior performance by vScreenML for this particular target. Targets seen by rfscore_VS during training of this method are marked with black triangles. **(B)** PPI benchmark, comprised of 10 protein targets. For each target, a single active complex is hidden amongst 2000 decoy complexes. Instead of using enrichment, the rank of the active compound (relative to the decoys) is calculated: thus, *lower* numbers indicate better performance. When comparing vScreenML against another method, a point *above* the diagonal indicates superior performance by vScreenML for this particular target. p-values in both cases were computed using the two-tailed Wilcoxon Signed-Rank test.

To evaluate in a statistically rigorous way which method was superior, we applied the (non- parametric) Wilcoxon Signed-Rank test: this paired difference test uses the rank values in the data, and thus it takes into account not just which method has higher EF-1%, but also the magnitude of the difference [63]. We used a two-tailed test, in order to assume no *a priori* expectation about what method would out-perform the other. At a threshold of p<0.05, this analysis shows that vScreenML out- performed 8 of the 9 alternate scoring functions to a statistically significant degree. Only RF-Score-VS was not out-performed by vScreenML at a statistically significant threshold; however, we note that about half of the 23 targets in this benchmark were included in training RF-Score-VS (black points in this figure), which may have provided it with a slight advantage relative to vScreenML (since the latter had not seen any of these targets before).

To test these methods on a second independent virtual screening benchmark, we drew from our own prior studies of inhibitors of protein-protein interactions [63]. In the course of evaluating existing scoring functions, we had several years ago assembled a set of small molecules that engage protein interaction sites; 10 of these protein targets had not been included in training vScreenML. For each of these, we had previously compiled 2000 decoys with dissimilar chemical structure matched to the active compound’s lipophilicity. The decoy compounds were already docked and energy minimized from our studies, making this “PPI set” a natural testbed for the newer methods that were not available at the time this benchmark was developed [63]. In contrast to the DEKOIS benchmark, the structures of the active complexes are drawn from (energy-minimized) crystal structures, removing a potential source of variability (since mis-docked active compounds should not be labeled “correct” by a scoring function).

Because each protein target is only associated with a single active compound in this test set, we cannot meaningfully calculate enrichment factor; instead, after scoring each of the complexes we simply report the rank of the active compound. As there are 2001 complexes for each protein target, a method that performs as random would be expected to rank the active compound at position 1001, on average. After applying each of the same scoring functions used in our DEKOIS experiment, we find that for 5 of the 10 protein targets vScreenML ranks the active compound among the top 100 (i.e., top 5% of the compounds for a given target) (**Figure 3b**). The other scoring functions tested each ranked the active compound in the top 100 for at most one target, except for RF-Score-VS which met this criterion twice. Once again applying the Wilcoxon Signed-Rank test to these rankings, we once again conclude that vScreenML out-performs at a statistically significance degree all of these alternate scoring functions except for RF-Score-VS.

**Figure s4:**
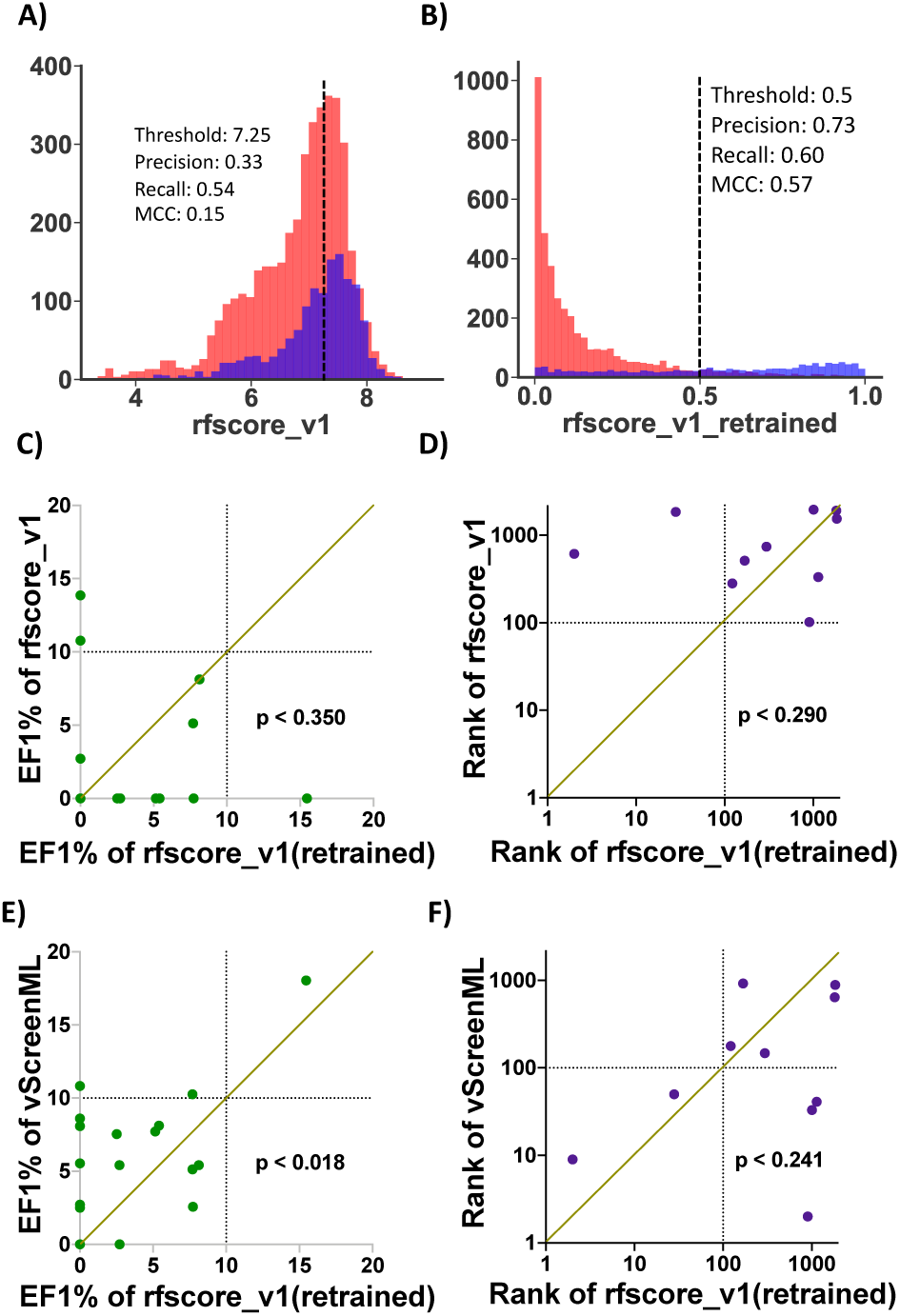
Retraining rfscore_v1 using D-COID. **(A)** Overlaid histograms for scores obtained when scoring active complexes (*blue*) and decoy complexes (*red*) from D-COID using the original rfscore_v1. **(B)** Overlaid histograms after re-training rfscore_v1. **(C)** Comparison of the original and re-weighted versions of rfscore_v1 applied to the DEKOIS benchmark. **(D)** Comparison of the original and reweighted versions of rfscore_v1 applied to the PPI benchmark. **(E)** Comparison of re-weighted rfscore_v1 versus vScreenML on the DEKOIS benchmark. **(F)** Comparison of re-weighted rfscore_v1 versus vScreenML on the PPI benchmark.

To determine whether vScreenML’s impressive performance derived from its training on the D-COID set or from the broad collection of features it includes, we used D-COID to train a model using the features from RF-Score v1; our re-trained model preserves the same random forest framework and hyperparameters from the original model [31]. As noted earlier (**Figure 1b**), RF-Score v1 initially yields very little discriminative power when applied to the D-COID set; after re-training on this set, we find much improved separation of the scores assigned to active versus decoy complexes (Figure S4ab), though not close to the performance of vScreenML (**Figure 2f**). This re-trained variant of RF-Score v1 also out-performs the original RF-Score v1 on both the DEKOIS and the PPI benchmarks, albeit not to a level of statistical significance, and for the PPI benchmark it even ranks two actives in the top 100 for their corresponding protein targets (Figure S4cd). That said, the level of improvement is insufficient for the re-trained RF-Score v1 to out-perform vScreenML in either benchmark (Figure S4ef), consistent with their relative performance on D-COID set. Overall, these observations show that training using the D-COID approach can certainly improve performance of existing scoring functions for other unrelated tasks; however, it also suggests that some part of vScreenML’s power derives from the broad and diverse set of features that it uses.

### Evaluating vScreenML in a prospective experiment

As noted earlier, it is absolutely essential to test new scoring functions in prospective experiments: this can readily determine whether performance in a given benchmark experiment is likely to extend into real future applications, and rule out any possibility that inadvertent information leakage allowed an overfit model to “cheat” in benchmark experiments. We selected as our representative target human acetylcholinesterase (AChE) because of its biomedical relevance and the availability of a straightforward functional assay (using commercially-available enzyme and substrate).

To ensure that our search for new candidate AChE inhibitors would not be limited by the chemical space present in a small screening library, we turned to a newly-available virtual library of “readily-accessible” but never-before-synthesized compounds [9]. At the time of our screen, this library was comprised of 732 million chemical entities that conform to historic criteria for drug-likeness [73,74]. Because building conformers and docking each entry in this library would be extremely computationally demanding, we instead took a two-step approach to finding candidate inhibitors. First, we explicitly docked a chemically-diverse set of 15 million representatives from the library, and applied energy minimization to the top 20,000 models from the crude docking step. We ranked each of these using vScreenML, and identified the top 100 candidates. For each of these 100 initial candidates, we returned to the complete compound library and identified 209 analogs on the basis of chemical similarity: after merging these with the parent compounds from each search, this led to a new focused library of 20,213 unique compounds. We structurally aligned each of these compounds back onto the parent docked model that led to their selection, re-minimized, and then used vScreenML to rank these second-stage candidates. We collected into a single list the 20 top-scoring compounds from the first round together with the 20 top-scoring compounds from the second round, noting that 4 compounds were included on both lists. We eliminated compounds that were extremely close analogs of one another, and sought to purchase the remainder. Based on a standard filter [75], none of these structures were predicted to be PAINS (pan-assay interference) compounds. Ultimately 23 compounds were successfully synthesized, as selected by vScreenML without any human intervention.

We initially tested these compounds at a concentration of 50 µM for inhibition of AChE, using a colorimetric enzyme assay (**Figure 4a**). To our amazement, we found that nearly all of the 23 compounds selected by vScreenML showed detectable enzyme inhibition: all except AC12 and AC7 showed a statistically significant difference in AChE activity relative to DMSO alone (p<0.05, one-tailed t-test). Of these 23 compounds, 10 of them provided more than 50% inhibition, indicating that these compounds’ IC_50_ was better than 50 µM. Moreover, the most potent of these used a variety of diverse chemical scaffolds, although the most potent pair (AC6 and AC3) do share an extensive common substructure (**Figure 4b**). We then evaluated the activity of the most potent inhibitor, AC6: in the absence of any medicinal chemistry optimization, we found this compound to have an IC_50_ of 280 nM, corresponding to a K_i_ value of 173 nM (**Figure 4c**). Thus, applying vScreenML led to a much higher hit rate than observed in typical screening campaigns, and also yielded a much more potent starting point than is typically observed.

**Figure 4:**
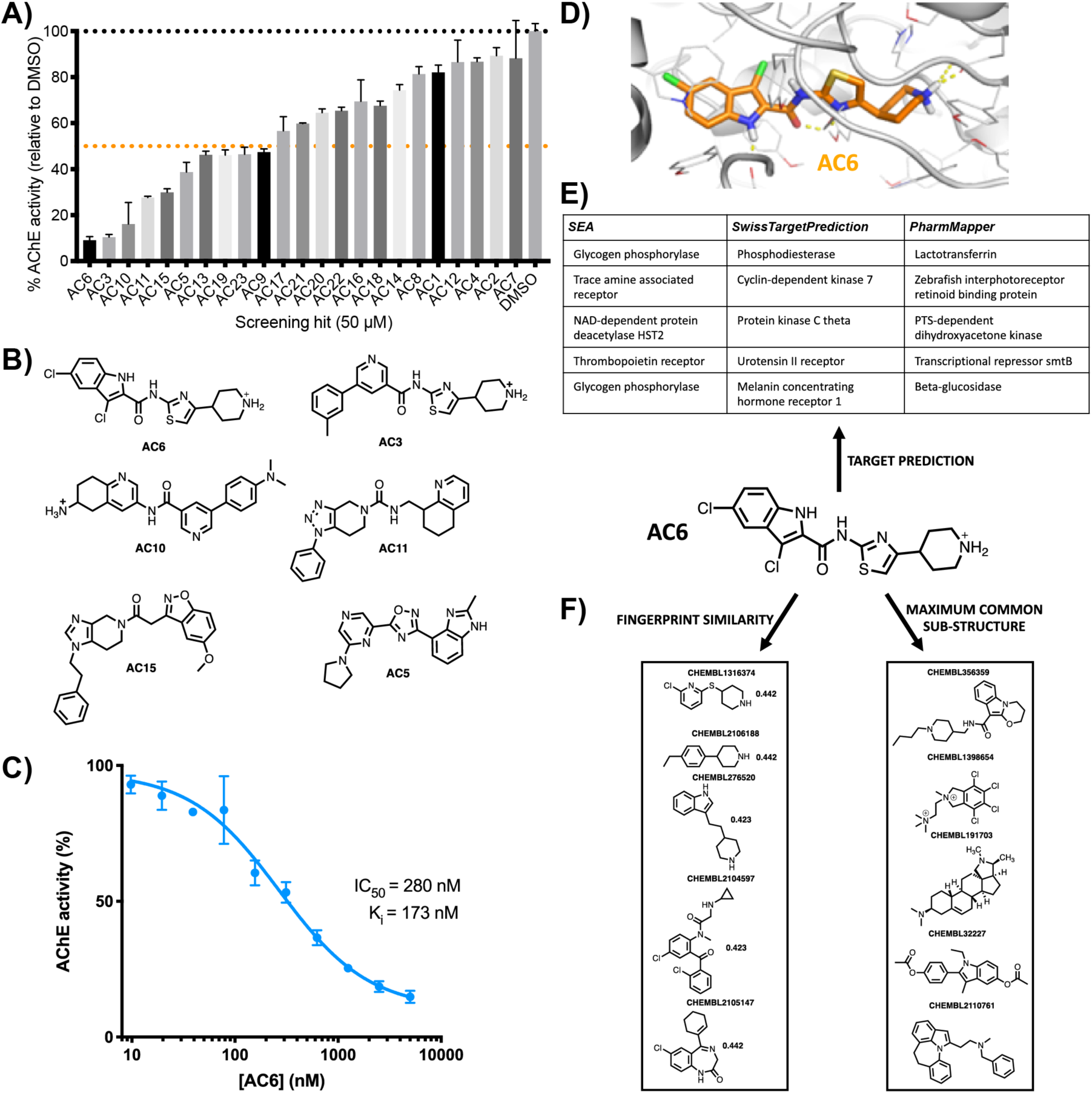
Prospective evaluation of vScreenML in a virtual screen against human acetylcholinesterase (AChE). **(A)** Of the 23 compounds prioritized by vScreenML for testing, at 50 μM nearly all of these inhibit AChE. Data are presented as mean ± SEM, n = 3. **(B)** Chemical structures of the most potent hit compounds. **(C)** Dose-response curve for the most potent hit compound, AC6. Data are presented as mean ± SEM, n = 3. **(D)** Model of AC6 (*orange sticks*) in the active site of the acetylcholinesterase (*light gray*). **(E)** Predicted activity of AC6 from three target identification tools: none of these identify AChE as a potential target of this compound, suggesting that this is a new scaffold for AChE inhibition. **(F)** Similarity searching against all compounds in ChEMBL designated as AChE inhibitors (either by fingerprint similarity of by shared substructure) finds no hits with discernible similarity, confirming that this is a new scaffold for AChE inhibition.

Unsurprisingly, the underlying model of the complex that was used by vScreenML to identify this compound shows extensive and nearly optimal protein-ligand interactions (**Figure 4d**). In principle, it should be the quality of these interactions that guided vScreenML to prioritize this compound for experimental validation. To rule out the possibility that vScreenML had instead somehow “recognized” AC6 as an AChE inhibitor from its training, we asked whether chemoinformatic approaches could have been used to find AC6.

We first provided the chemical structure of AC6 to three different “reverse screening” methods: Similarity Ensemble Approach (SEA) [76], SwissTargetPrediction [77,78], and PharmMapper [79,80]. Each of these tools look for similarity of the query compound against all compounds with known bioactivity, then they rely on the fact that similar compounds have similar bioactivity to predict the likely target(s) of the query compound. SEA and SwissTargetPrediction carry out this search on the basis of 2D similarity (i.e. similar chemical structures), whereas PharmMapper evaluates 3D similarity (i.e. shared pharmacophores). We took for each method the top 5 predicted activities for AC6, but found that none of these methods included AChE among their predictions (**Figure 4e**). All of these methods do include AChE among their list of potential targets, however, as confirmed by ensuring that this prediction emerges when these servers are provided with the structure of previously-described AChE inhibitor donepezil (**Figure S5**).

To directly determine the AChE inhibitor described to date that is most similar to AC6, we compiled from ChEMBL all 2742 compounds reported to have this activity. We then screened this collection to determine their similarity to AC6, as defined by either chemical fingerprints or by shared substructure. The 5 most similar compounds as gauged by either approach bear no obvious similarity to AC6 (**Figure 4f**): collectively then, these experiments confirm that AC6 is indeed a novel chemical scaffold with respect to its inhibition of AChE, and could not possibly have been identified by vScreenML through inadvertent leakage during the model’s training.

**Figure S5.**
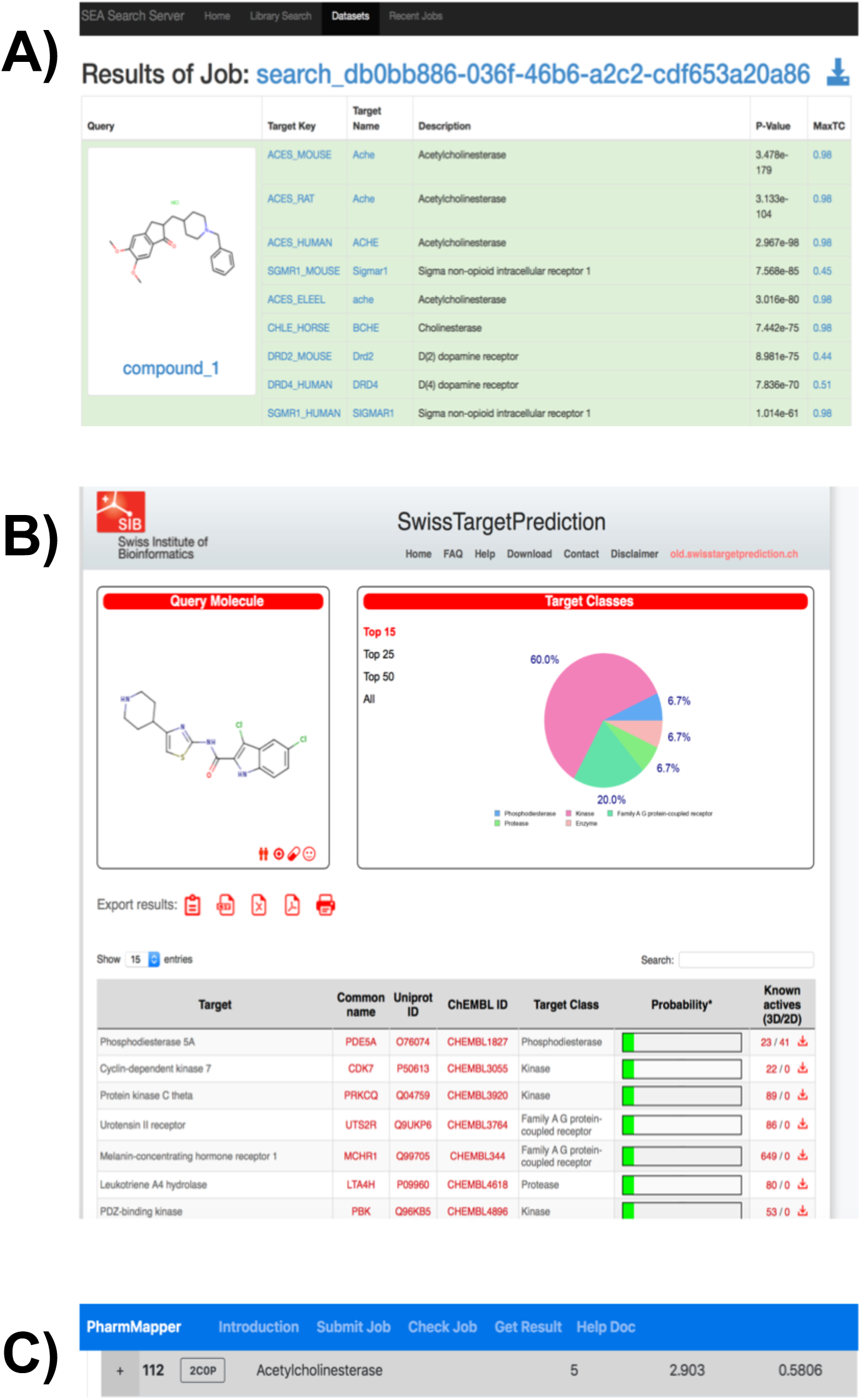
Positive control for target identification methods. We confirmed that all three methods would successfully identify AChE as the target of a known AChE inhibitor (donepezil, CHEMBL1678). **(A)** Similarity Ensemble Approach (SEA). **(B)** SwissTargetPrediction. **(C)** PharmMapper. We note that AChE was only ranked #112 among the PharmMapper hits because the 3D conformations it built for donepezil were not sufficiently well-matched to the active conformation to produce a better ranking.

## Discussion

At the outset of this work, we noted that typical virtual screening studies report hit rates of about 12%, with the most potent reported compound having K_d_ or K_i_ value of ∼3 µM (with the caveat that some of these relied on additional optimization beyond the initial screen) [12]. Obviously the results of our screen against AChE using vScreenML far surpass these mileposts; in light of this, it important to carefully consider the potential contributions to vScreenML’s performance in this experiment.

First, we re-emphasize the dissimilarity between AC6 and any known AChE inhibitor: this makes it exceedingly unlikely that vScreenML found AC6 simply on the basis of having been trained on some close analog.

Second, we carried out a non-standard two-step screening strategy to efficiently explore the complete Enamine collection, hoping to essentially carry out an internal round of medicinal chemistry optimization before testing any compounds explicitly. Tracking the provenance of our most potent compounds, however, we discovered that all four of our most potent compounds had already been identified in the first of the two screening steps (**Table S1**). A previous virtual screen of the Enamine library [9] explicitly docked all compounds from the library, at a time that the library comprised “only” 138 million compounds, and found through retrospective analysis that picking a single representative compounds from a cluster of analogs would typically not yield sufficient docking score for the cluster to be advanced for further exploration. In essence, both our results and the observations from this previous screen suggest that the SAR landscape may not be sufficiently smooth to allow potentially promising scaffolds to be identified from a single arbitrary representative: rather, finding the best hits (on the basis of docking scores) does unfortunately require explicitly screening each member of the library individually. In this context, then, it is unlikely that the observed performance of vScreenML can be attributed to having used a two-step strategy for screening the Enamine library.

**Table s1:** Provenance of AChE inhibitors. For each of the 10 AChE inhibitors that provided more than 50% inhibition at a concentration of 50 μM, we determined at what stage this compound was prioritized for testing. Our strategy included two stages of screening: first we screened only 15 million diverse compounds from the Enamine collection, then we expanded our search by collecting analogs for each of these hits. We note that 7 of these 10 compounds were identified in the first round of screening; after re-refinement in the second round, 3 of these were still highly-ranked whereas 4 had been surpassed by analogs (or received lower scores upon re-refinement). Only 3 of these 10 compounds would have been missed if our screening had been limited to a single round of 15 million compounds.

| Compound | Identified as a top-scoring hit |
| --- | --- |
| AC6 | 1<sup>st</sup> round |
| AC3 | 1<sup>st</sup> round |
| AC10 | Both rounds |
| AC11 | 1<sup>st</sup> round |
| AC15 | 2<sup>nd</sup> round |
| AC5 | 2<sup>nd</sup> round |
| AC13 | 2<sup>nd</sup> round |
| AC19 | Both rounds |
| AC23 | 1<sup>st</sup> round |
| AC9 | Both rounds |

In this vein, we also note that our screening strategy was allowed to explore an unusually large chemical space comprising 732 million synthetically-accessible compounds. However, seven of our top ten compounds (those with IC_50_ values better than 50 µM) had already been identified in the first screening step (**Table S1**), owing to the ineffectiveness of identifying useful scaffolds from a single representative compound. The bulk of the success in this screen was essentially achieved by screening a library of 15 million diverse compounds, which is by no means unprecedented and has not led to such dramatic success in the past.

Importantly, we cannot rule out the prospect that the performance we observe here is a result of AChE being an unexpectedly easy target. It is certainly the case that virtual screening hit rates against GPCRs are often much higher those obtained for other target classes [12]. Indeed, careful examination of the literature showed that some of the studies reporting virtual screens against AChE [81-85] do indeed find considerably higher hit rates and more potent compounds than the median values we quote across all target classes. In light of these other results, then, a degree of caution must be exercised before extrapolating the performance of vScreenML in this prospective AChE benchmark to other target classes; further evaluation will be needed to explicitly determine whether vScreenML affords similarly outstanding results in future screening experiments.

At the same time, however, results of retrospective benchmarks comparing vScreenML to other scoring functions are unambiguous. As described, vScreenML dramatically outperforms eight other modern machine learning scoring functions on both the DEKOIS and the PPI benchmark sets. Both benchmarks were carried out with careful vigilance to ensure that information from training could not contaminate the test data. In the past, we strongly suspect inadvertent overtraining of this type has limited the utility of other models and at the same time provided artificially inflated performance on initial (retrospective) benchmarks. Indeed, a recurrent disappointment from many past machine learning scoring functions has been their inability to translate performance from retrospective benchmarks into equivalent results in future prospective applications [37]. For example, three years after publication of nnscore [32] this program was used in a screen against farnesyl diphosphate synthase, and only provided one hit with IC_50_ of 109 μM (from ten compounds tested) [86]. Where possible, then, we strongly urge incorporation of careful prospective evaluations alongside retrospective benchmarks, as a safeguard against potentially misleading performance from the latter. Already such prospective experiments have been included in other recent studies [41,87], strongly supporting transferability of the underlying methods. The ability to readily compare vScreenML against other machine learning scoring functions was also greatly facilitated by the Open Drug Discovery Toolkit (ODDT) [88], which provides implementations of multiple methods. Direct head-to-head evaluations of this manner are indeed critical to explore the relative strengths of different approaches, ideally across diverse types of benchmarks.

While vScreenML does incorporate a broad and distinct set of features, these have been largely collected from other approaches: there is nothing particularly unique or special about the features it includes. There are also numerous potential contributions to protein-ligand interactions that are not captured in this collection of features, ranging from inclusion of tightly-bound interfacial waters [16,89,90] to explicit polarizability and quantum effects [91,92]. In this vein, ongoing research in ligand-based screening has led to new approaches that learn optimal molecular descriptors (and thus the representation that directly leads to the features themselves) at the same time as the model itself is trained [93,94]: these might similarly be used as a means to improve the descriptors used in structure-based screening as well. Thus, there is likely to be considerable future improvement to vScreenML that is possible, by further optimization of the features that it captures.

Rather than the specific features incorporated in this first incarnation of vScreenML, we believe that the impressive performance we observed in our retrospective benchmarks is instead primarily attributable to the strategy used in training the model. Whereas scoring functions have historically focused on recapitulating binding affinities of complexes, vScreenML is unique in having been trained to distinguish active complexes from extremely challenging decoys in the D-COID set. Indeed, the overarching hypothesis of our study was that building truly compelling decoys to better represent the (inactive) compounds selected from actual virtual screens we would lead to a model capable of distinguishing precisely these cases. The performance of vScreenML in both the retrospective and prospective benchmark strongly supports this hypothesis.

Thus, the D-COID set represents an important resource for driving development of improved scoring functions beyond vScreenML, and accordingly we have made this dataset freely available for this purpose (see *Methods*).

## Methods

### Accessing these tools

The D-COID dataset is available at https://data.mendeley.com/datasets/8czn4rxz68/ [95]. vScreenML is available at https://github.com/karanicolaslab/vScreenML .

### Building the D-COID set

The overarching goal of our study was to train a model for real virtual screening applications. We therefore included in D-COID only active complexes that included representative drug-like ligands, and excluded chemical matter that did not reflect the composition of the screening libraries we prefer to use.

We downloaded from the Protein Data Bank (PDB) [96] all protein-ligand complexes (56,195 entries as of May 2018), and then restricted this set to crystal structures with resolution better than 2.5 Å (43,148 complexes). We then drew from Ligand Expo [97] to define a set of 2937 specific ligands found in the PDB that we deemed ineligible for our study: these include nucleotide-like molecules (e.g., ATP), co-factors (e.g., NAD), metal-containing ligands (e.g., heme), crystallographic additives (e.g., PEG), and covalent ligands. We filtered to retain only complexes that included an eligible ligand, and did not have an additional ligand within 12 Å of the eligible ligand (leaving 26,271 complexes). To focus training on precisely the type of chemical matter used in our virtual screens, we then applied to this collection the same stringent filter we use when building our screening libraries: molecular weight between 300-400 Da and clogP 1-4. This filter drastically cut down the size of our collection (to 2,075 complexes). Finally, complexes with double occupancy or ambiguous density were manually excluded, leaving a high-quality collection of 1,383 active complexes.

For each of these active complexes, we extracted the ligand and used the Database of Useful Decoys Enhanced (DUD-E) [56] server to generate 50 property-matched decoys: compounds with similar physicochemical properties but dissimilar chemical topology. For each of these decoy compounds, we used OpenEye’s OMEGA [98] to generate 300 low-energy conformers, and then used ROCS [57] to align each of these to the structure of the active conformer from the PDB. The three decoys that best matched the three-dimensional shape and pharmacophoric features of the active conformer were identified on the basis of their Tanimoto-Combo score; this led to a total of 4,149 decoy compounds. By virtue of having aligned the conformers of the decoys to the active conformation to evaluate their similarity, already the alignment was available for placing the decoy compound in the corresponding protein’s active site. We later discovered that 39 of these decoy compounds included chemical features that could not be processed by the programs we used to extract structural features for vScreenML; these decoys were removed, leading to a total of 4,110 decoy complexes.

Finally, to present both the active and decoy complexes in a context mimicking that of a virtual screening output, we subjected all complexes to standard energy minimization in Rosetta [61].

### Extracting structural features

For each of the minimized active and decoy complexes, structural features were extracted first using the Rosetta (“REF15”) energy function [61]. Ligand properties were calculated using ChemAxon’s cxcalc [69], and the ligand’s conformational entropy was estimated using OpenEye’s SZYBKI tool [70]. The open source implementations of RF-Score [31] and BINANA [68] were used to calculate structural features from these two programs. The complete list of vScreenML’s features is presented in **Figure S1**.

### Machine learning

We considered a total of nine classification algorithms in this study, using the Python implementations of each: Support Vector Machine (SVM) [99], Gradient Boosting (GB) [100], Extreme Gradient Boosting (XGB) [101], Random Forest (RF) [102], Extremely Randomized Trees (ET) [103], Gaussian Naïve Bayes (GNB) [104], k-Nearest Neighbor (kNN) [104], Linear Discriminant Analysis (LDA) [104], and Quadratic Discriminant Analysis (QDA) [104].

Training was carried out using 10-fold cross-validation; splitting the dataset into 10 subsets was carried out in a stratified manner to ensure that the overall ratio of actives to decoys was preserved in each split. For XGBoost hyperparameter optimization, we carried out a grid search to find the set of parameters that gave the best cross-validation accuracy (splitting out a separate validation set from the data).

To re-train RF-Score v1 under D-COID, we used a standard random forest model with hyperparameters n_estimators=500 and max_features=5 (drawing these values from the original study describing RF-Score v1 [31]).

### Virtual screening benchmarks

Comparisons between scoring functions was enabled by the Open Drug Discovery Toolkit (ODDT) [88], which provides implementations of nnscore (version 2), RF-Score v1, RF-Score v2, RF-Score v3, PLEClinear, PLECnn and PLECrf at https://github.com/oddt/oddt. The implementation of RF-Score-VS was obtained from https://github.com/oddt/rfscorevs.

In both the DEKOIS and the PPI benchmark experiments, we carefully sought to minimize any potential information leakage from vScreenML’s training (on D-COID) and the targets present in these benchmark sets. Excluding a specific complex present in both sets is insufficient, because the structure of a close chemical analog bound to the same target protein could still provide an unfair advantage. For this reason, we excluded from these benchmarks sets any protein targets present in D-COID (on the basis of shared Uniprot IDs). This reduced the number of DEKOIS targets from 81 to 23, and the number of PPI targets from 18 to 10.

For the DEKOIS set, we docked both the actives and the decoys to their respective target protein using OpenEye’s FRED [64], then applied energy minimization in Rosetta. For the PPI set, active complexes were minimized starting from their crystal structures; decoy complexes were generated by docking with FRED then energy minimized.

Statistical analysis was carried out using the (two-tailed) Wilcoxon Signed-Rank test as implemented in Python. Comparisons were applied directly to the EF-1% values for the DEKOIS experiment, and to the log10 of the ranks in the PPI experiment.

### Virtual screen against acetylcholinesterase

We began by downloading from the chemical vendor Enamine the “diverse set” of 15 million compounds representative of their REAL database (732 million compounds). For each compound we used OMEGA [98] to generate 300 low-energy conformers, and used FRED [64] to dock these against the crystal structure of human acetylcholinesterase solved in complex with potent inhibitor donepezil (PDB ID 4ey7) [105]. We carried forward the top 20,000 complexes (on the basis of FRED score) for Rosetta minimization, and used each of these minimized models as input for vScreenML.

For each of the top 100 complexes (as ranked by vScreenML), we extracted the ligand and used this to query the Enamine database for analogs. Each query returned 210 analogs; because 787 of these were redundant, this led to a new collection of 20,213 unique compounds for the second stage of screening. Each of the compounds in this new library was used to build 300 conformers, and ROCS was used to select the conformer that allowed for optimal alignment onto the ligand in the complex from the first round of screening. The resulting models were energy minimized in Rosetta, then used as input for vScreenML.

Models from both the first and second rounds of screening were collected together, and the top-ranked models from vScreenML were identified, and the top-scoring 32 compounds were requested for synthesis. Of the requested compounds, 23 were successfully synthesized and delivered for testing.

### Acetylcholinesterase inhibition assay

Compounds were tested for inhibition of human acetylcholinesterase (AChE) using a colorimetric assay [106]. Acetylthiocholine is provided as substrate, which is hydrolyzed by AChE to thiocholine; the free sulfhydryl then reacts with Ellman’s reagent (5,5’-dithiobis-(2-nitrobenzoic acid); DTNB) to yield a yellow product that we detected spectrophotometrically at 410 nm. AChE, acetylthiocholine, and DTNB were acquired together as the Amplite^TM^ Colorimetric assay kit (AAT Bioquest). Assays were carried out in 0.1 M sodium phosphate Buffer (pH 7.4), 1% DMSO, with 0.01% Triton. Assays were carried out in 96-well plates in reaction volumes of 100 µL, and absorbance was monitored for 30 min. The rate of product formation was determined by taking the slope of the absorbance as a function of time, and normalized to that of DMSO alone to yield percent inhibition for each well.

IC_50_ values were obtained from dose-response curves spanning inhibitor concentrations from 10 nM to 50 µM. To determine K_i_, we first determined the K_m_ value for substrate acetylthiocholine under our assay conditions. This allowed the Cheng-Prusoff equation [107] to be used for obtaining K_i_ from IC_50_, assuming classic competitive inhibition.

### Novelty of AC6 as an AChE inhibitor

For each of the target identification methods (Similarity Ensemble Approach (SEA) [76], SwissTargetPrediction [77,78], and PharmMapper [79,80]), we used the corresponding web servers to generate predictions for AC6.

To find the most similar known AChE ligands, we searched ChEMBL [108] for acetylcholinesterase and downloaded all 2742 hits in SDF format. We then used ChemAxon’s Standardizer tool to remove counterions from compounds in salt form. Using RDKit [109] we generated Morgan fingerprints with radius of 2 for each of the ChEMBL ligands, then evaluated the Dice similarity of these fingerprints relative to that of AC6. We also used RDKit to evaluate the maximum common substructure (MCS) between AC6 and each of the ChEMBL ligands, setting ringMatchesRingOnly=True and completeRingsOnly=True. We ranked the resulting matches based on the number of atoms and bonds in the common substructure.

## Acknowledgements

We thank Joanna Slusky for a useful suggestion regarding presentation of the figures, and Juan Manuel Perez Bertoldi for his initial application of vScreenML to the PPI benchmark set. We thank ChemAxon for providing an academic research license. This work used the Extreme Science and Engineering Discovery Environment (XSEDE) allocation MCB130049, which is supported by National Science Foundation grant number ACI-1548562. This work was supported by grants from the National Institute of General Medical Sciences (R01GM099959, R01GM112736, and R01GM123336) and from the National Science Foundation (CHE-1836950). This research was funded in part through the NIH/NCI Cancer Center Support Grant P30 CA006927.

## Notes

https://data.mendeley.com/datasets/8czn4rxz68

